# Long term context dependent genetic adaptation of the viral genetic cloud

**DOI:** 10.1101/262345

**Authors:** Tzipi Klein, Antonio V. Bordería, Cyril Barbezange, Marco Vignuzzi, Yoram Louzoun

**Author notes:** Corresponding author (YL).

## Abstract

RNA viruses generate a cloud of genetic variants within each host. This cloud contains high frequency genotypes, and a very large number of rare variants. While the dynamics of frequent variants are affected by the fitness of each variant, the rare variants cloud is affected by more complex genetic factors, including context dependent mutations. It serves as a spearhead for the viral population’s movement within the adaptive landscape. We here use an experimental evolution system to show that the genetic cloud surrounding the Coxsackie virus master sequence slowly, but steadily, evolves over hundreds of generations. The evolution of the rare variants cloud often precedes the appearance of high frequency variants. The rare variants cloud’s evolution is driven by a combination of a context-dependent mutation pattern and selection for and against specific nucleotide compositions.This combination affects the mutated dinucleotide distribution, and eventually leads to a non-uniform dinucleotide distribution in the main viral sequence. We then tested these conclusions on other RNA viruses with similar conclusions.

## INTRODUCTION

**Some of the most difficult diseases to treat or cure are caused by RNA viruses.** The global AIDS pandemic and the repeating seasonal flu epidemics are both examples of diseases caused by RNA viruses (1–3). One of the main obstacles in the treatment and vaccination against such diseases is the particularly high mutation rate of RNA viruses (10^−3^ to 10^−5^ substitutions per nucleotide copied (4–6)). This mutation rate induces a rapid adaptation to the changing environment, and the resulting development of resistance to various vaccines and drugs (7). In addition to high mutation rates, RNA viruses also present short generation times, and large population sizes, as the number of viral particles in an infected organism can be higher than 10^12^ virions (8). The combination of high mutation rates, large population sizes, and short generation times results in a complex and dynamic mutant distribution termed viral quasispecies (9), viral cloud or mutant spectrum. The concept was first developed by Eigen, Schuster, and Biebricher to describe error-prone replication and self-organization of primitive macromolecules thought to carry information as precursors of more complex life forms (10–13). It was later found relevant in the understanding of RNA virus dynamics, as the viral mutant distribution undergoes a continuous process of genetic change and selection (14).

**The composition of viral sequences within a single host contains a high frequency master sequence (often the consensus sequence) and a mutant spectrum - the viral “cloud”.** Due to the high mutation rate and population size of RNA viruses, theoretically even a single virion infection can quickly evolve into a collection of related viral genomes, containing every possible point mutation and many double mutations (15). As a result it would only take a few rounds of replication for the viral cloud to expand in sequence space and lose important biological information, if not for the continuous selection against unfit variants, and viral robustness mechanisms (16). This combination of mutation and selection determines the composition of the viral cloud. Eigen proposed that selection not only works on each specific unfit variant, but that the quasispecies cloud as a whole also acts as a unit of selection (10–13). Whether this is the case for RNA viruses remains to be determined; some evidence suggests that, at least under certain growth conditions, selection of a group of minority variants may occur (17–19).

**The analysis of viral genetic variants, due to technical limitations, has mostly focused on the evolution of a specific subset of minority variants bearing phenotypic (fitness) alterations, such as amino acid substitutions.** However, with the advent of Next Generation Sequencing (NGS) it is possible to enlarge the analysis beyond a limited set of variants with macroscopic frequencies, and expand the study to low frequency variants that were previously undetectable (20–22). We here describe the dynamics of these rare variants. While these rare variants are short-lived, the full cloud evolves over the scale of hundreds of generations. While selection of the main minority variants is determined by their specific fitness, the rare variants cloud is affected by a genetic selection mechanism (i.e. selection mechanism not determined by the proteins produced).

**Viral nucleotide composition is affected by the host environment.** In particular, patterns of codon usage are strongly correlated with overall genomic GC content (23). Viruses also avoid specific nucleotide pairs, such as CpG, in order to mimic their hosts’ CpG usage and avoid detection by the host (24–26). CpG repression is also affected by codon position, with repression being much weaker between codons (25). APOBEC, a host antiviral protein, also affects the A to G mutation composition (27). Finally, viral mutations have been shown to maintain a constant GC-content (28,29). Virus consensus dinucleotide abundance is also affected by the host, mostly due to CpG repression (30). For example, viruses transferred from avian to human hosts have been shown to change over decades (31–34). Here we show that at the short time scale within a single host, the dinucleotide composition of mutant variants derived from the main sequence significantly differs from that of the consensus sequence and that this difference is the result of a context-dependent mutation pattern. Our results demonstrate several important points regarding the population structure of viruses: 1) The viral population is composed of high frequency variants and a rare variants cloud, 2) the cloud of mutants is genetically linked, 3) the cloud evolves over hundreds of viral life cycles. 4) The high frequency variants follow a regular selection process, while the rare variants cloud moves ahead, following complex dynamics based on context-dependent mutation patterns and genetic selection.

## MATERIAL AND METHODS

### Viral sequences

NGS data from a previous experiment (19) was used. Three different variant of CVB3 virus were used: Wild type variant, low fidelity variants and high fidelity variant. Each independent passage started from a single isolate of each of the CVB3 virus variants, and passages were carried out in triplicate for forty passages (representing around 120 viral generations). The infections were kept until full cytopathic effect (CPE) was observed, around 24 hours, before preparing the new passage. To prepare the next passage, cultures were freeze-thaw, centrifuged to obtain a clean viral supernanant and a new passaged was prepared by diluting the obtained supernanant to have a new infection at a multiplicity of infection of 0.01. Deep sequencing was done for passages 1, 3, 5, 7, 9, 11, 13, 15, 17, 19, 20, 21, 23, 25, 27, 29, 31, 33, 35, 37, 39, and 40, and the first and last passages were resequenced as technical replicates (19).

To verify our results in other viruses, NGS data from two variants of NDV were used. 1) Newcastle disease vaccine (B1 type, LaSota strain). 2) An NDV natural infection, obtained from a chicken coop at Klahim moshav in southern Israel. The viral genome was sequenced using an Illumina sequencer. In addition, we used influenza virus NGS data. Wild-type influenza A virus (A/Paris/2590/2009 (H1N1pdm09)) was passaged 5 times (at least 15-20 replication cycles) in triplicate on MDCK cell monolayers in 6-well plates. Cells were infected at an MOI = 0.001 and 48 hours post-infection, influenza A viruses were harvested in clarified supernatant and virus titres (TCID50 or plaque assay) were determined at each passage. Viral RNA genome was extracted from supernatants (Macherey-Nagel), reverse transcribed with Accuscript High Fidelity 1st strand cDNA Synthesis kit (Agilent) using 5’-AGCRAAAGCAGG-3’ primer, and amplified by PCR using Phusion High-Fidelity DNA Polymerase (ThermoScientific). Eight PCRs were designed to cover the coding regions of the eight genomic segments (primer sequences are available upon request). The PCR products were fragmented (Fragmentase), multiplexed, clustered, sequenced in the same lane with Illumina cBot and GAIIX technology.

### Quality control

Initial quality control of the fastq files was performed using FastQC v0.10.1(35). Since the quality of the edges of the reads was inconsistent, six nucleotides from each edge were trimmed. Reads that had more than twenty percent nucleotides with a quality score of less than twenty were removed. The remaining reads were mapped to the known viral genome using bowtie v1.0.0. (36). Reads with a mapping score lower than fifty, and over expressed reads were removed. The clean files were again tested using FastQC v0.10.1 and samples that did not show good results on all standard indices were removed. In addition, all samples from passage one were removed, as they did not have enough time to develop a full cloud. Only the open reading frame (ORF) was used.

### Data organization

The mapping of the reads to the known viral genome was performed using bowtie v1.0.0. The resulting sam files were used to calculate the frequencies of nucleotides, dinucleotides and nucleotides triplets in every position over the viral genome. These frequencies were used during downstream nucleotide composition analysis.

### Frequent variants reconstruction

High frequency minority variants full sequences were reconstructed using QuasiRecomb-1.2(37) with default parameters. This reconstruction was done separately for each sample and gene. Aligned sam files from previous steps were used for this analysis.

### Variants mutation frequency

For each read, the genetic distance from the consensus sequence of the sample was calculated. These distances were used to determine the frequency of mutations from the consensus over time. To estimate the same frequency in high frequency minority variants full sequences, those reconstructed sequences were used. The genetic distance was calculated as the number of mutations of each such variant from the most common variant in the sample. A variant is considered shared if it appears in samples from two or more independent experiments (different virus variant or duplicate).

### Statistical analysis

Poisson λ parameter for the fit to the frequency of mutations per read was estimated as the average number of mutation from the consensus sequence per read. The value of λ was than divided by read length to estimate mutation rate per nucleotide. This λ parameter was used to calculate the theoretical poisson distribution.

Correlations were calculated using spearman’s rank correlation coefficient.

Wilcoxon signed-rank test was used to test the significance of some of our results. Both one sided and two sided tests were used. In the case of two sided test, paired measures from the same sample were used.

Analysis of variance (ANOVA) was used to confirm the significance of the effect of neighbouring nucleotides on the mutation profile. ANOVA was calculated separately for each specific mutation (e.g. C to A) and Bonferroni correction was used. The range of p values is shown.

### Information theory

To estimate the genetic diversity of each sample, Shannon Entropy was used. Entropy can be written as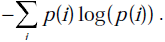 We used the number of possibilities as the base of the log. The entropy of the clouds was calculated per position using nucleotide frequencies obtained from all the sequenced reads that passed the quality control. The entropy of the high-frequency variants was calculated per gene sample using the frequencies of the reconstructed genes.

### Calculating nucleotides and dinucleotides frequencies and mutation rates

Nucleotides and dinucleotides frequencies were directly calculated as the average frequencies in the nucleotide and dinucleotide frequency tables. Frequencies of mutations were calculated as the frequencies after removing the nucleotides at the consensus from the tables for each position in sample. Mutation rates were estimated as nucleotides frequencies by nucleotides in the consensus. For example the mutation rate from A to G was calculated as the frequency of G only at positions that had A in the consensus. The Mutation rate from A to G with a 3’ C was calculated as the frequency of G only at positions that had an A and a C before at the consensus. The mutation rate was then normalized by the average mutation rate for each specific mutation, to leave only the effect of neighbouring nucleotides. The neighbour affected mutation rate of high-frequency mutations was calculated, for example, as frequency the of mutations from A to G with a 3’ C out of all the mutations from A to G, and later normalized by the average frequency of mutations from A to G, to fit the scale of the low-frequency mutation rates.

For calculations of the expected dinucleotide mutation rates, the frequencies of dinucleotides in the consensus sequence and the mutation rates from one nucleotide to another were used.

### Rare variant cloud, synonymous and non-synonymous computation

For each sample, all positions with entropy higher than the average entropy by more than ten standard deviations were found. During rare variants cloud calculations, these positions were ignored, to neutralize the effect of the high-frequency variants.

For synonymous calculations, we removed all mutations in the nucleotide frequency tables that resulted in amino acid change compared to the consensus. Positions that had no possible mutations as a result were ignored. The same was done for mutations that did not cause an amino acid replacement in non-synonymous calculations.

### Estimating the rate of cloud evolution

The per position nucleotide Shannon entropy was used. We normalized the entropy by nucleotide and codon position, by calculating the average entropy of all consensus nucleotides in each codon position. We than divided the entropy of each position by the fitting average entropy. This calculation was performed for all samples. For each experiment, the normalized entropy of samples from the fifth, seventh and ninth passages were correlated to that of later samples. The changes in those correlations following different lags (number of passages between samples) were used as an estimate of the rate of cloud evolution.

### Calculating entropy correlations

Spearman’s rank correlation coefficient was calculated between nucleotide frequency Shannon entropy for each position in the genome, and same entropy for positions k nucleotides away in each sample. To account for codon position, the correlations were calculated separately between first and second codon positions, third codon position and first codon position at the following codon, etc. The same calculation was performed for entropy normalized by codon position and nucleotide entropy, and for entropy normalized by codon position, nucleotide and first neighbour nucleotides in the consensus.

## RESULTS

### The viral population is multilayered with different within host mutation depths

**We studied NGS data of three different variants of Coxsackievirus B3 (CVB3): wild type variant (WT), low fidelity variant (S299T) and high-fidelity variant (A372V).** Each variant was independently evolved in triplicate for forty passages, starting from a single isolate, with each passage representing 2-3 viral generations giving us a total of around 120 generations. MOI was kept at 0.01 for all the 40 passages minimizing the effect of bottlenecks. Every odd passage, the twentieth passage and the last passage were deep sequenced. Technical replicates of the first and last passages were re-sequenced for quality control. The average coverage per sample was 10,968-fold (Table S1). The experimental procedures are detailed in a previous publication (19). Briefly, 70-nucleotide reads were sequenced, and aligned to the viral consensus sequence. We then produced full reads of the main minority variants in each gene using the QuasiRecomb-1.2 Software (37) (see Methods). Variant frequency can be estimated locally at the single read level, or at the full-gene level. Note that full genome reconstructed variants may not be reliable and are thus not analysed here.

**Most observed mutations are real and not sequencing errors.** The variant frequency at the read level was calculated using every read that passed the quality control (Fig S1), without filtering for statistically significant mutations. Due to the sequencing and methodological errors mixed with this data It was not used to study specific mutations but overall trends, as the methodological error should affect all samples in a similar way. To validate that most of observed mutations are not methodological errors (PCR, sequencing…), and can be used to study the properties of the genetic cloud, we calculated the percentage of mutations in the cloud over time by virus variant (Fig S2A). As expected, the mutation percentage rose over time, as mutations accumulated (r = 0.85, p-value = 1.15e-52 for spearman correlation coefficient). WT variant accumulated fewer mutations than the low fidelity variant and more mutations than the high-fidelity variant (Two-sided Wilcoxon signed-rank test p-value < 0.05 for both WT and low fidelity variant, and WT and high-fidelity variant). The mutation percentage over time was also calculated by codon position. The mutation percentage was significantly different between codon positions (Two-sided Wilcoxon signed-rank test p-value < 9.013e-07 between codon positions). This result is unlikely to be the result of sequencing errors, but rather by the codon composition and synonymous-nonsynonymous mutation difference. After combining those two effects (Fig S2B), by the end of the analysis, the low fidelity virus at third codon positions had about four times the initial mutation load of the high-fidelity virus at the first codon position. Thus, at least in the at last period of the experimental evolution a high enough (at least 75 %, but probably much more) portion of the mutations in our data are not the results of sequencing errors and can be used to study the genetic cloud. Since there were no significant differences in the biases obtained in the early and later periods of the experiment, we conclude that during most of the experiment, the vast majority of observed mutations are not artefacts. Note that the signal obtained is much stronger than a difference of 25 %. Thus, even in the extreme unlikely case that all the mutations in the early period are sequencing errors, our results are still valid.

**Highly mutated variants appear more than expected randomly.** The mutation frequency was calculated as the fraction of mutations per read compared to the consensus sequence of the sample. The variant distribution did indeed follow a Poisson distribution up to a mutation probability of 2.e-3 per nucleotide per sequencing passage (Fig 1A) (see Methods). Beyond this point, the distribution did not fit a Poisson any more, showing significantly more highly mutated variants than expected randomly. This deviation shows that the neutral process by itself cannot explain our observations, and that other mechanisms, such as positive selection or biased mutation profiles (i.e. not all reads have equal mutation rates) must be considered. When translating the computed nucleotide mutation rate to the gene level, an estimated 34%-93% of sampled sequences of each gene carried at least one mutation (except for gene 3B that is exceptionally short, 65 Nt). Some of these mutations were high-frequency variants that repeatedly appeared in different replicates of the experiment, but the large majority represented rare variants (Fig 1B).

**A clear difference was observed between the high frequency minority variants, and the large cloud of rare variants.** While the frequent variants appeared at some stage and then stabilized, with divergent variants taking over the population (Fig 1B), the rare variant frequency slowly, but steadily, increased over time even before the more frequent ones emerged, as can be observed from the regression coefficient of the mutant frequencies vs. time (Fig 1C and 1D). The cumulative frequency of the rare variant was at least 99.91 % of the total viral population and cannot be neglected. Thus, as will be further shown, the cloud of rare variants preceded the emergence of the more frequent ones and may have facilitated their appearance.

**Fig 1.**
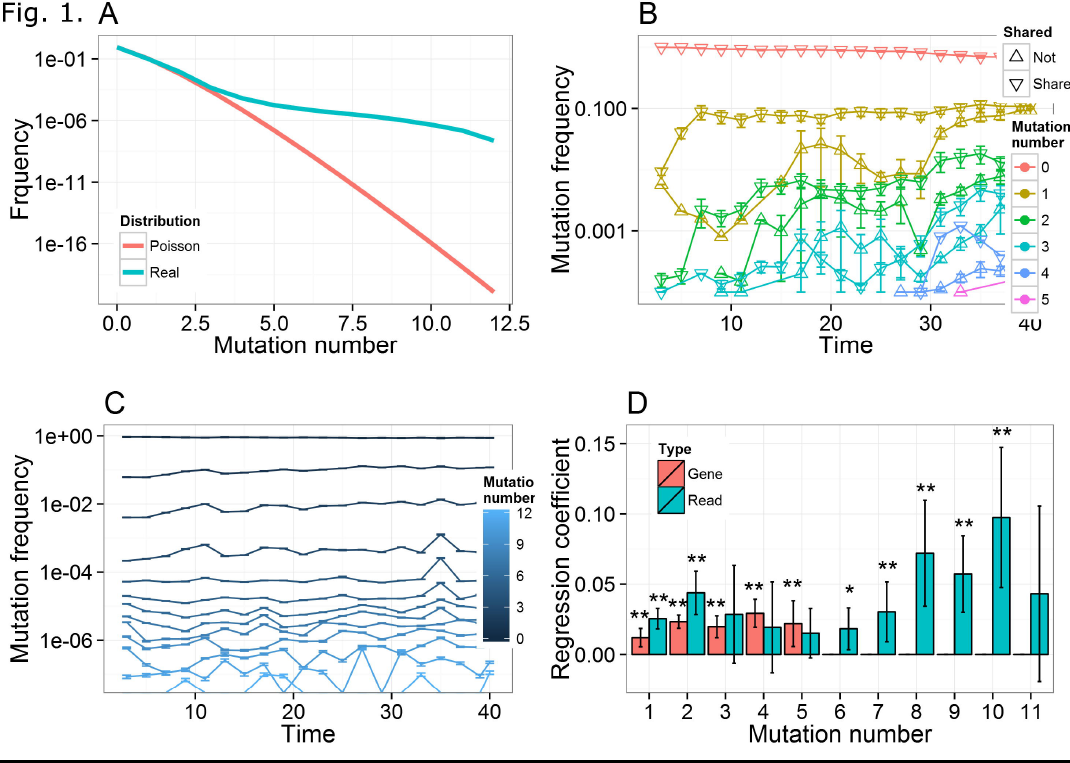
Mutation number per read and gene. The frequency of mutations from the consensus sequence for each sample was calculated per read and reconstructed gene (See Methods). (A) Fit of the frequency of mutation number over all reads to Poisson distribution. (B) Frequency of mutation number per reconstructed gene over time. Error bars depict standard error over genes. A variant that appears in more than one independent experiment is considered shared. (C) Frequency of mutations per read over time. Error bars depict standard error over reads. (D) Regression coefficient of mutation number per read and reconstructed gene by time. Values were normalized by average frequency. Error bars depict 95% confident intervals. ** represents p-value <0.01, and * represents p-value < 0.05

**Our results could theoretically be explained as selection for specific high frequency mutations, or protein level functionality specific to this virus.** To prove that this is not the case, the same calculations were performed considering only synonymous mutations (see Methods) with similar results, as shall be further shown. **To validate that the observed synonymous mutations are not sequencing errors, we tested the effect of time on the synonymous mutations rate.** The percentage of synonymous mutations rose over time (r = 0.837, p. value = 1.4132e-48 for spearman correlation coefficient), and were strongly affected by variant fidelity (Two-sided Wilcoxon signed-rank test p-value < 0.05 for both WT and low fidelity variant, and WT and high fidelity variant) (Fig S2C), showing that the observed mutations are not sequencing artefacts.

### The rare variants cloud evolves slowly but steadily over many generations

**Many of the frequent minority variants taking over the population were shared among experiments and are argued to be selected through a higher fitness of the resulting viral phenotype (Fig 1B)(19, 38).** However, the rare variants cloud may determine the availability and initial frequency of different variants for the selection process. To estimate the importance of the rare variants cloud, we computed the per position nucleotide distribution entropy (see Methods). The entropy represents the deviation from the consensus at each position (Fig 2A). Higher entropy implies a larger fraction of the population in the mutation cloud. The entropy calculations were highly reproducible over the technical replicates, further validating this method (Fig S3) (average r over all replicates = 0.7713, standard error = 0.0401 (n=17), p-value < 4.8333e-168 for Spearman correlation of nucleotide entropy of all technical replicates). The average entropy increased along the experiments, until it reached a maximal value and then stabilized (Fig 2B) (r = 0.88, p-value = 5.46e-60 for spearman correlation against time). This behaviour was similar over all sub-strains studied and over all repeats of the analysis.

**Fig 2.**
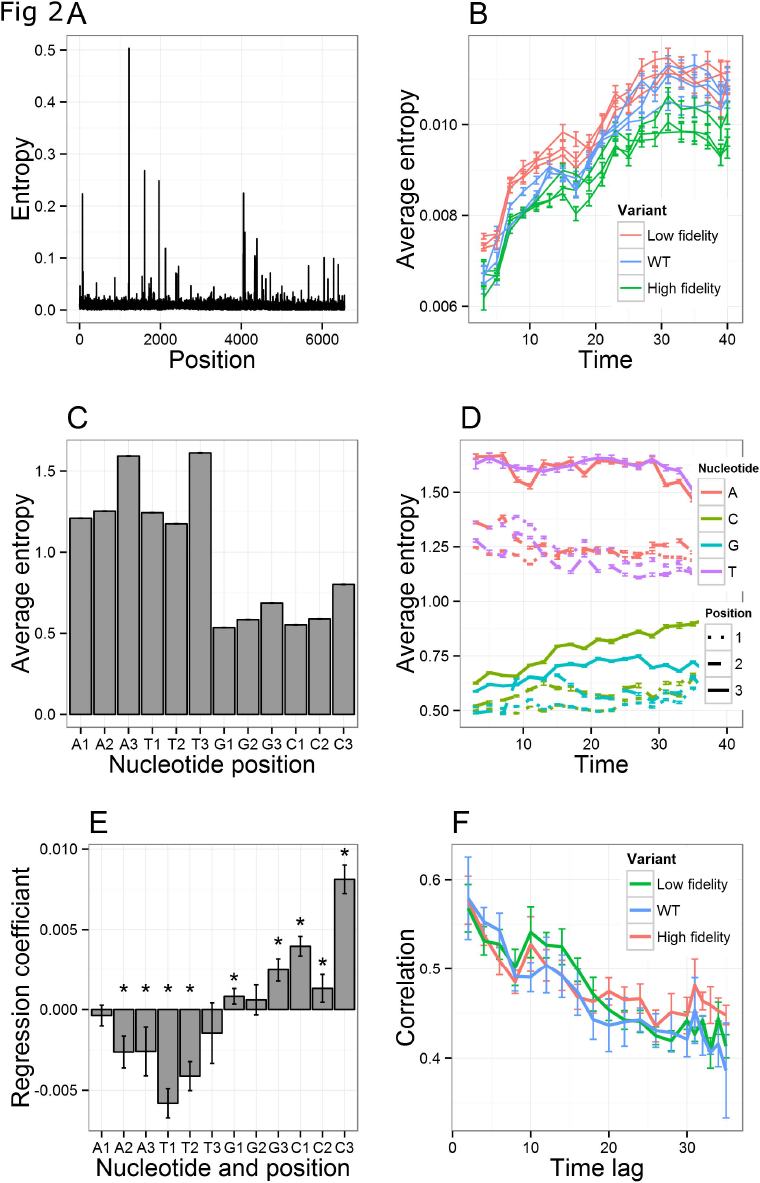
The viral cloud moves slowly but steadily over many generations. (A) Nucleotide entropy per position. One sample was used as an example (wild type duplicate 1, passage 15). (B) Average entropy per sample over time. Error bars depict standard error. Results were smoothed over a window of 5. (C) Average nucleotide entropy by nucleotide and codon position. Error bars depict standard error. (D) Average nucleotide entropy over time by nucleotide and codon position. Error bars depict standard error. (E) Regression coefficient of average nucleotide entropy by nucleotide and codon position vs. time. Error bars depict 95% confident intervals. (F) Correlations of per position nucleotide entropy between samples from the same experiment, at time t and time t + lag. Entropy is normalized by average entropy based on the consensus sequence nucleotide and codon position. Error bars depict standard error. * p-value< 0.01

**The per-position entropy was strongly affected by the codon composition, with a clear difference between the first two positions in each codon and the third position in the codon (Fig 2C) (Two-sided Wilcoxon signed-rank test p-value < 1e-30).** An even stronger difference was observed between positions with A and T, compared with positions with C and G in the consensus (Fig 2C, D) (Two-sided Wilcoxon signed-rank test p-value < 1e-30). While the latter difference could be explained by the stability of the molecular bonds, the first could be the effect of selection, since third position mutations are typically synonymous mutations. Indeed, when computing the entropy of synonymous mutations, the most significant difference was between AT and CG in the third position (Fig S4C) (Two-sided Wilcoxon signed-rank test p-value < 1e-30). All other results were conserved when studying only synonymous mutations (Fig S4), showing that the evolution of the cloud is not solely directed by amino-acid based selection.

To limit the entropy to the rare variants cloud, positions where the entropy was higher than the average entropy by more than ten standard deviations were removed (Fig S5). These positions typically represent the high frequency variants. The results were similar to the results for the whole population, indicating that the results above were caused by the rare variants cloud and not the high frequency variants. The entropy of each position followed approximately the average entropy over the entire sequence.

**To understand the evolution of specific nucleotides in the cloud, we divided the entropy per position in the nucleotide by the average entropy of the sample.** We found that the normalized entropy evolved very slowly, but consistently (Fig 2D). The entropy of Cytosine slowly increased over time, while the entropy of Adenine and Thymine decreased. The strongest changes were the slow increase of Cytosine in third position (C3), and the decrease in the entropy of Thymine in first position (T1) (Fig 2E).

*The rare variants cloud starts evolving before the main frequent variants.*

**The slow changes in entropy show that the rare variants cloud evolves at a slow pace.** To produce a quantitative estimate of the change rate, we computed the cross-correlation between the entropy profile (entropy as a function of position along the sequence) between each time **t** and time **t + lag.** The effects of codon positions and the difference between AT and CG clearly induced correlations in different times. However, even when the entropy in each position was normalized by the average entropy as a function of the nucleotide and the position in the codon (see Methods), the correlation remained high for a long time. The correlation dropped by a factor of approximately 0.15 over 30 passages (r = −0.67, p-value = 2.97e-59 for spearman correlation) and would expectedly decrease to 0 over approximately 120 passages (each passage is 2-3 generations), showing a very slow evolution of the genetic cloud over a period of 240-360 viral generations (Fig 2F).

**The same result held when only synonymous mutations were considered, and when high-frequency variants were removed from the analysis (Fig S4F and S5F), showing again that the observed phenomena are a generic process affecting the rare variants cloud.** Interestingly, the entropy of the high-frequency variants rose slower than the rest of the cloud (Fig S6 and 2B), further supporting the concept that the rare variants cloud may drive the emergence of the high-frequency variants, and could serve as a predictor for the emergence of future high frequency variants.

### The rare variants cloud variants are genetically linked

**To understand the dynamics of the rare variants cloud, we first analyzed its genetic structure.** The simplest characteristic of the cloud is the nucleotide distribution, which was slightly biased toward A in the consensus and toward G in the mutants, following a high A to G mutation frequency (Fig 3A – inset). The distribution became more interesting when the frequency of dinucleotides was computed (Fig 3A). The dinucleotide frequencies were biased, and cannot be explained by the single nucleotide frequencies. This bias has been speculated to be the result of dinucleotide stacking energies and properties of cell environments(39), such as CG repression(24,25).

**Fig 3.**
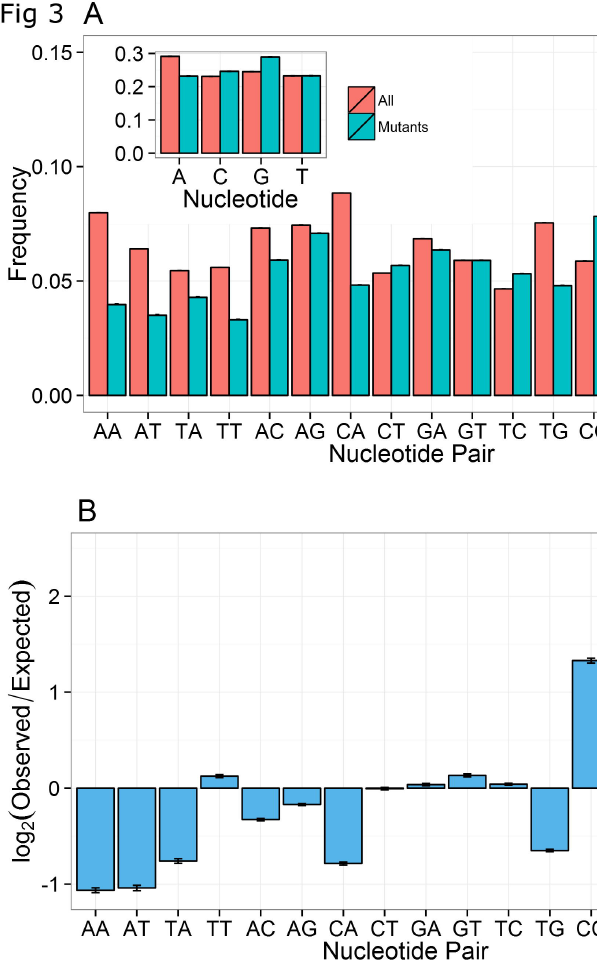
Dinucleotides mutation frequencies are biased. (A) Dinucleotides frequencies in the full cloud, and only for mutations. Dinucleotides that are different from the consensus were considered mutations. Error bars depict standard error. Same analysis was performed for single nucleotides in the insert. (B) Log of the ratio between observed and expected frequency of mutant dinucleotides for each dinucleotide (see Methods). Error bars depict standard error.

**The mutation cloud surrounding the consensus sequence had a completely different distribution (Fig 3A).** The frequency of all dinucleotides differing from the consensus sequence (non-consensus dinucleotide – NCDN) was computed and found to differ from the distribution in the consensus in all dinucleotide pairs (Fig3A) (Two-sided Wilcoxon signed-rank test p-value < 1e-30 for all dinucleotide pairs except GT). One could consider the following model for the observed NCDN distribution. If the entire cloud was produced from random mutations, the probability of observing dinucleotide CT (for example) can be computed as the sum over all possible combinations in the consensus sequence multiplied by the mutation probabilities. For example:

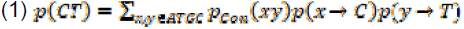

**All elements of the model can be computed from the observed consensus and variants.** The observed distribution was very different from the expected one using the model above (Fig 3B) (one-sided Wilcoxon signed-rank test p-value < 1e-12 for all dinucleotide except TC, CT and GA). One possible explanation for the difference could be that mutations in neighbouring positions are correlated. However, double mutations (mutation in neighbouring positions in the same read) amount to less than 0.94% of the cloud. A simpler explanation would be the effect of the high-frequency portion of the variants taking over. However, those variants typically carry 1-2 mutations per gene (Fig 1B) and do not affect the rare variants cloud.

**Another possible way to explain the divergence from the expected distribution is that specific dinucleotides were preferentially produced, maintained or lost.** In the following sections, we will show that this is indeed the case. Such a result suggests either a strong genetic based selection or a mechanism of induced mutations, either by the host or by viral molecules.

### Preferential generation of dinucleotides – Effect of neighbouring positions

**The peculiar structure of the NCDN cloud structure can be understood through the correlation between the entropy of neighbouring nucleotides.**The correlation was computed between the entropy in each position and the entropy in positions k nucleotides away. The entropy in first and second neighbour nucleotides was correlated along all positions in the codon (Fig 4A) (One-Sample Wilcoxon signed-rank test p-value < 1e-14 for first and second neighbours at all codon positions). Surprisingly, the correlation was negative in first neighbours and positive in second neighbours (One-Sample Wilcoxon signed-rank test p-value < 1e-20 for distances 1-4). The entropy correlations were constant over time (Fig 4B). The negative correlation can be explained by the different probability of fixation of different nucleotides and the dinucleotide bias. There are fewer CG pairs in the consensus than other dinucleotides, and AT have higher entropy than CG. Since AT were more often than randomly expected near CG, a negative correlation between the entropy in nearest neighbours emerged. Indeed, when the entropy in each position was divided by the expected entropy based on the consensus sequence nucleotide and codon position, the negative correlations disappeared. However, surprisingly, the correlation at distance 2 was still much higher than the correlation between neighbouring positions or positions 3 nucleotides away (Fig 4C) (two-Sample Wilcoxon signed-rank test p-value < 1e-30). This was not a codon effect, since the different correlation was consistent over all positions in the codon, as well as nucleotides in different codons (Fig 4A) (One-Sample Wilcoxon signed-rank test p-value < 1e-14 for first and second neighbours at all codon positions). If the correlation would have been induced by the presence of conserved regions, the normalized correlation would be maximal at a distance of 1.

**Fig 4.**
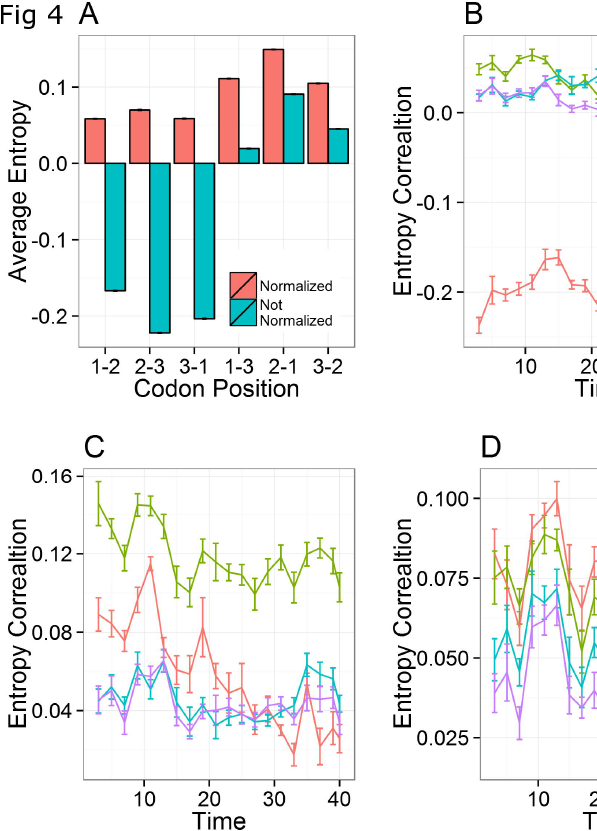
Neighboring nucleotides entropy correlations. The correlation between the entropy in each position and the entropy in positions k nucleotides away was computed for all samples. Error bars depict standard error. (A) First and Second neighbors Nt along all codon positions. (B) Entropy correlations over time. (C) Normalized entropy correlations over time. The entropy was normalized by the average entropy based on the consensus sequence nucleotide and codon position. (D) Normalized entropy correlations over time. The entropy was normalized by the average entropy based on the consensus sequence nucleotide, its first neighbors and codon position.

**A simple, yet surprising explanation for this correlation was that mutations were affected by the nucleotide in neighboring positions.** In other words, mutations near an A nucleotide would be rarer than mutations near a C nucleotide.

In such a case, for example:

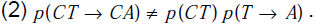

**The outcome of such a context-dependent mutation profile would be correlations in the entropy in nucleotides at distances of 2 and 3.** Since AT and CG have different entropies, then the codon usage could induce correlations between distant positions. Indeed, when entropy was normalized by the effect of neighbouring positions (see Methods), the correlation decreased with distance between nucleotides (Fig 4D).

**To confirm that the mutation profile was affected by neighbouring positions, we computed the probability of mutation from each nucleotide to each nucleotide, as a function of the 5’ and 3’ neighbouring nucleotide (Fig 5A).** Surprisingly, both the 3’ and 5’ neighbouring positions affected the probability to mutate (Fig 5A) (ANOVA 1.88e-305 ≤ p-value ≤ 0.16 for neighbour nucleotides). The biggest difference in the mutation rate was between different mutations, mostly between transitions and transversions. The effect of neighbouring nucleotides remains even after normalizing for specific mutations (e.g. C to A) (Fig 5B) (see Methods). The variance of the normalized specific mutation frequencies, as effected by their 5’ and 3’ neighbouring nucleotides, rose over time (Fig S7) (spearman correlation r=0.535, p-value = 4.69e-14 for variance over time), further suggesting that this is not purely the effect of biased sequencing and methodological errors.

**Fig 5.**
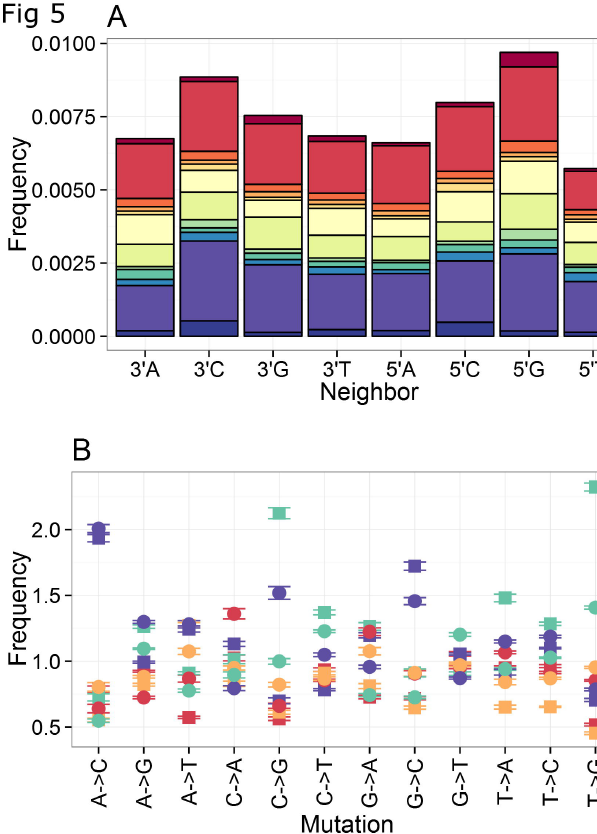
Mutation profile is affected by neighboring positions. Probabilities of mutations from one nucleotide to another, as a function of the 5’ and 3’ neighboring nucleotides were calculated. Error bars depict standard error. (A) Cumulative frequencies. (B) Normalized by specific mutation.

**This effect could in principle result from the codon structure.** We thus repeated the analysis for synonymous and non-synonymous mutations (Fig S8). The results were similar for non-synonymous mutations (ANOVA 1.51e-290 ≤ p-value ≤ 4.86e-7 for neighbour nucleotide). For synonymous mutations the effect was weaker but still present (ANOVA 0 ≤ p-value ≤ 0.58 for neighbour nucleotides), indicating that the mutation profile was affected by codon structure, but not fully explained by it. The analysis was also performed at each codon position by itself (e.g. all nucleotides at position 1 of the codon etc.) (Fig S9) with similar results. The biased mutation probability explained the dinucleotide structure and the correlation, but more interestingly, it may affect the dinucleotide distribution of the consensus.

**To further examine the observed position dependent mutation probability, we studied the mutation profile of high frequency mutations.** High frequency mutation frequencies were calculated as the frequencies of mutations in the reconstructed genes (see methods), while accounting for neighbour nucleotides and mutation type. Frequencies were normalized by the average mutation type frequency to fit the scale of low-frequency mutations. The high frequency mutation profile was affected by neighbouring nucleotides (Fig S10A), and the effect was similar to that of the low-frequency mutations (Fig 5B, S10A, S10B) (Spearman correlation for high frequency mutations, and low frequency mutations r = 0.5452579, p. value = 9.237e-09). This result further supports the possibility of context-dependent mutation rate affecting nucleotide distribution at all levels, from low-frequency mutations to high-frequency mutations, and possibly even the consensus dinucleotide distribution.

**Such context-dependent mutation rate was previously studied in other organisms, such as eukaryotes, yeast, bacteria, and bacteriophages (40–45).** Various mutation-causing mechanisms were suggested to be affected by neighbouring nucleotides, causing this effect in mutation rates. These mechanisms include polymerase fidelity and proofreading, mismatch repair, mismatch stability and Dcm methylation (45–47). With the exception of polymerase fidelity, all of these mechanisms affect DNA. Thus, they and are not relevant here, as they cannot affect viral single stranded RNA.

**The current results suggest that the genetic dynamics affecting the rare variants cloud are not simply a combination of random mutations and purifying selection.** Instead, the rare variants cloud is driven by a context-dependent mutation mechanism and complex selection patterns.

**This claim emerges from results obtained from experimentally evolved populations.** To check that this may be a generic feature of RNA viruses, we analysed NGS data from three additional RNA viruses: Newcastle disease virus (NDV) vaccine, natural infection of NDV and Influenza virus. As was the case for the Coxsackie virus experimental samples, the mutation profile of those viruses was affected by neighbouring nucleotides (Fig 6) (ANOVA 1.10e-10 ≤ p-value ≤ 5.07 for neighbour nucleotide for Influenza virus, where we had enough repeated experiments).

**Fig 6.**
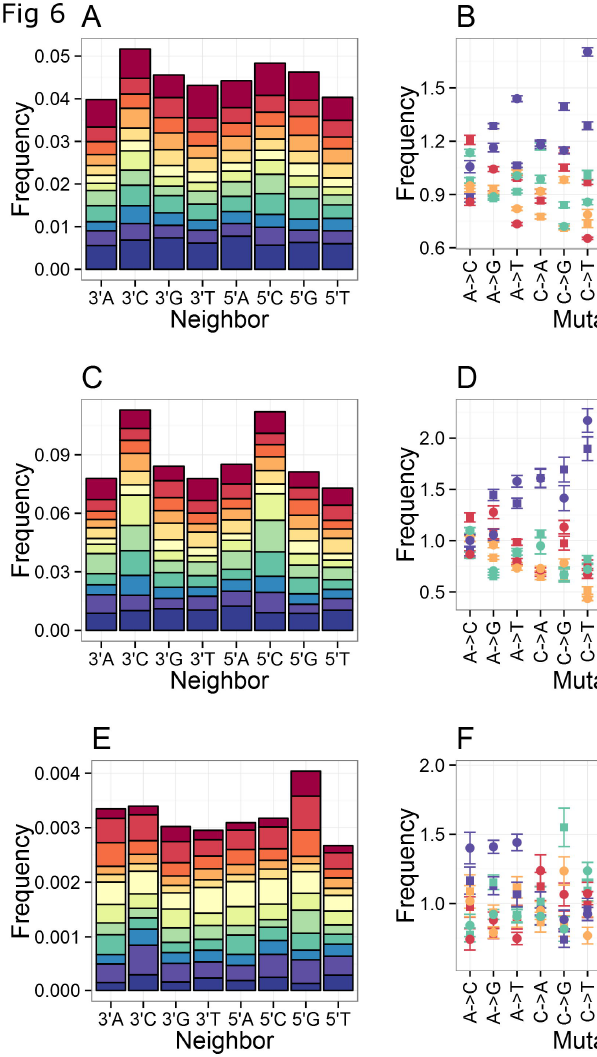
Mutation profile is affected by neighbouring positions in Newcastle disease virus. The same calculations as in figure 5 were performed for NGS data from: (A,B) Newcastle disease virus vaccine (LaSota) (C,D) natural infection of Newcastle disease virus. (E,F) Influenza virus. For Newcastle disease vaccine and Newcastle disease virus natural infection error bars depict standard error within the sample, as there is only one sample each.

## DISCUSSION

**Virus populations constitute a cloud of genetic variants.** This cloud is often composed of a dominant genotype, a few high-frequency minority variants and a very large number of rare variants. While the dynamics of the dominant genotype and high-frequency variants are easier to monitor and have been the focus of research, the dynamics of the rare variants cloud that encompasses most of the observed sequences has not been fully studied yet. Here, we studied the dynamics of this cloud and shown in multiple strains and samples of Coxsackie virus that it slowly evolves over hundreds of replication cycles. This slow evolution precedes the selection and emergence of higher frequency minority variants and may be the background supporting their emergence.

**The main data analysed in this study comes from a longitudinal evolutionary study in which a homogeneous population of Coxsackie virus B3 was passaged for 120 generations to a new, never encountered before, cell type, A549.** During these passages, we found that mutations in the whole viral population do not follow a Poisson distribution, where rare outliers are more frequent than expected from the Poisson distribution.

**The cloud can be described through a bilayered distribution of variants in the viral population: a layer composed of higher frequency shared variants and another broader layer composed of rare variants.**

**The lack of fit to Poisson distribution forfeits the possibility of neutral local evolution and implicates that either strong positive selection or another mechanism was acting.** One salient observation was that some of the high frequency variants in the population appeared at some point and increased to dominate the population. These variants depict a more classical positive selection based on the fitness advantages of beneficial mutations. Indeed, in a previous study on the adaptation of CVB3 to A549 cells, there was a strong selective process for beneficial mutations in the structural proteins of the virion, VP1, VP2 and VP3 involved in binding to the primary and secondary viral receptors (CAR and DAF) on the host cell. When we decoupled these high frequency variants from the rest of the mutations in the population, we unveiled the rare variants cloud layer, which according to our calculations evolved slowly over time, but precede the emergence of these variants. This steady increase in variability goes in concordance with a starting homogeneous population that generates new variability without any bottlenecks.

**Some of the mutations in the rare variants cloud can be attributed to the well-described lack of proofreading mechanisms in viral RNA-dependent RNA polymerases (48).** Indeed, our entropy results show that the lowest fidelity variant 3D-S299T replicates had higher average entropy compared to the other variants. It is possible that the observed context-dependent mutation rate is affected by a context-dependent bias in the viral RNA-dependent RNA polymerases activity. Such an explanation will be consistent with the fact that the context-dependent mutation profile seems to differ from one virus to another, as RNA viruses usually encode their own unique RNA-dependent RNA polymerase (49).

**Another possible explanation is that a significant fraction of mutations in viruses are induced, and not simply the result of random replication errors.** Such mutations may result from RNA editing, such as the ADAR and APOBEC families (50–53). ADAR and APOBEC are believed to be mechanisms of antiviral defense in the cell, mainly for retroviruses where this effect is well proven, although for single stranded RNA viruses this function is still contested (27,54–56). If this is the case, we could expect mutation patterns to be affected by the binding sites of these enzymes. However, proteins of the ADAR and APOBEC families can only cause A to G and C to U mutations, and as a result they cannot fully explain the observed context-dependent mutation profile. In addition, the mutation pattern was also affected in a natural infection of Newcastle disease virus in chicken. As the chicken APOBEC family is incapable of mediating C to U mutations (31,57), it cannot explain the mutation profile by itself.

**A further possibility is that the observed cloud is a result of either positive or negative selection acting upon this initial pool of variants.** This selection may be for structural stability (58) or GC content (28,29). If this is indeed the case, one could conclude that the entire cloud evolves, and rare variants are producing other rare variants. This could explain the large number of variants with many mutations, not expected if all mutations would originate from the consensus sequence.

**The genetic diversity of the cloud can be estimated by several methods (59).** An interesting observation resulting from our analysis of the genetic diversity of the rare variants cloud is the effect of the neighboring sequence on mutation. Here, we used nucleotide composition entropy to measure genetic diversity. Other methods such as the fraction of mutated sequences in each position yield similar results. Entropy is affected by nucleotide composition as well as codon position. Mutations at third codon position are more likely to accumulate in the population, as these mutations usually result in synonymous mutations (60). Adenine and Thymine are more likely to mutate than Cytosine and Guanine. As a result, nucleotide composition entropy is naturally biased, and should be accounted for. However, even when accounted for, the observed composition of the cloud could not be explained.

**We show here that cloud composition is the result of context-dependent mutation patterns and the probability of observing a given mutation (e.g. C to A) is determined by its neighboring nucleotides.** Strikingly, the observed effect of neighboring positions occurred regardless of codon position and thus, was not limited to within single codons. Moreover, the analysis of different viruses revealed similar patterns. Such patterns also appeared when only accounting for high-frequency mutations, indicating that this phenomenon could also affect the long term mutation accumulation in the consensus. Those mutations are mostly affected by selection, but a bias in the initial pool of available mutations could affect the mutation distribution. The context-dependent mutation rate, as well as the need to avoid certain dinucleotides (e.g. CG), induced correlations in the entropy profiles, as well as a highly biased dinucleotide composition. While this has been demonstrated here for four viruses, it may likely be true for others.

**Another important observation emerging from our analysis is that the rare variants cloud seems to move continuously toward a different dinucleotide composition than the main variant, rather than remaining stationary or moving at random.** The possible constraints governing directionality are not clear besides the observed genetic structure and effect of neighboring nucleotides giving preference to certain dinucleotides and not others. It is possible that weak positive selection is also acting on this component of the population. Nevertheless, this bilayered structure of the viral genetic variants cloud could have important implications on the evolvability of the population and predicting the emergence of higher frequency variants. Our data suggest that the rare variants cloud may be moving ahead of the most frequent sequence, and in a sense spearheading the evolution of the consensus sequence and the emergence of new Master sequences. We hypothesize that the direction taken by the rare variants cloud may influence which high frequency variants will be selected and fixated through classical selection processes in the whole population. The rare variants cloud selection could be due to maintain the aforementioned genetic structure but some not very strong positive selection might be action too. This hypothesis will be further tested in future research.

## DATA AVAILABILITY

The raw data used for this project can be found in the BioProject database under the accession identifier PRJNA420867.

## ACKNOWLEDGEMENT

We thank Saar Tal for preparing the Newcastle disease virus samples for sequencing.

## SUPPORTING INFORMATION

**S1Table. Details and coverage of all samples.**

*Number of samples that passed quality control appears in parenthesis (see Methods for further details).

**S1 Fig. Quality control indices before and after cleaning.** One sample was used as an example (wild type duplicate 2, passage 17). (A) Per base sequence quality before cleaning. (B) Per base sequence Nt content before cleaning. (C) Per sequence GC content distribution compared to theoretical normal distribution before cleaning. (D) Per base sequence quality after cleaning. (E) Per base sequence nucleotide content after cleaning. (F) Per sequence GC content distribution compared to theoretical normal distribution after cleaning.

**S2 Fig. Frequency of mutation in the cloud over time by virus variant.** (A) Full cloud (B) Full cloud by virus variant and codon position (C) Synonymous mutations cloud. Error bars depict standard error.

**S3 Fig. Nucleotide entropy is reproducible over technical replicates. Per** position nucleotide entropy of all samples of both technical replicates.

**S4 Fig. The viral synonymous mutations cloud moves slowly but steadily over many generations**. Identical to figure 2, only for synonymous mutations (see Methods).

**S5 Fig. The viral rare variants cloud moves slowly but steadily over many generations.** Identical to figure 2, only for rare variants cloud (see methods).

**S6 Fig. Entropy of high frequency variants rises slowly.** Average entropy of reconstructed high frequency variants over time per gene in sample. Error bars describe standard error.

**S7 Fig. Context dependent mutation variance rises over time.** The variance of the normalized specific mutation frequencies, as effected by their 5’ and 3’ neighbouring nucleotides over time. Error bars describe standard error.

**S8 Fig. Identical to figure 5, divided to synonymous and nonsynonymous mutations.** (A,B) Synonymous (C,D) Non-synonymous.

**S9 Fig. Identical to figure 5, divided by codon position.** (A,B) first codon position (C,D) second codon position and (E,F) third codon position.

**S10 Fig. High frequency mutation profile is affected by neighboring positions.** Neighbor nucleotides frequency was calculated per mutation from one nucleotide to another in the reconstructed genes, compared to the consensus sequence of each sample (see methods), as a function of the 5’ and 3’ neighboring nucleotides, and were normalized by specific mutation. (A) High frequency mutations by mutation type and neighbor. (B) High frequency normalized mutations plotted against low frequency normalized mutations.

